# Tafazzin regulates the function of lipopolysaccharide activated B lymphocytes in mice

**DOI:** 10.1101/2021.05.17.444507

**Authors:** Hana M. Zegallai, Ejlal Abu-El-Rub, Laura K. Cole, Jared Field, Edgard M. Mejia, Joseph W. Gordon, Aaron J. Marshall, Grant M. Hatch

## Abstract

B lymphocytes are responsible for humoral immunity and play a key role in the immune response. Optimal mitochondrial function is required to support B cell activity during activation. We examined how deficiency of tafazzin, a cardiolipin remodeling enzyme required for mitochondrial function, alters the metabolic activity of B cells and their response to activation by lipopolysaccharide in mice. B cells were isolated from 3 month old wild type or tafazzin knockdown mice and incubated for up to 72 h with lipopolysaccharide and cell proliferation, expression of cell surface markers, secretion of antibodies and chemokines, proteasome and immunoproteasome activities, and metabolic function determined. In addition, proteomic analysis was performed to identify altered levels of proteins involved in survival, immunogenic, proteasomal and mitochondrial processes. Compared to wild type lipopolysaccharide activated B cells, lipopolysaccharide activated tafazzin knockdown B cells exhibited significantly reduced proliferation, lowered expression of cluster of differentiation 86 and cluster of differentiation 69 surface markers, reduced secretion of immunoglobulin M antibody, reduced secretion of keratinocytes-derived chemokine and macrophage-inflammatory protein-2, reduced proteasome and immunoproteasome activities, and reduced mitochondrial respiration and glycolysis. Proteomic analysis revealed significant alterations in key protein targets that regulate cell survival, immunogenicity, proteasomal processing and mitochondrial function consistent with the findings of the above functional studies. The results indicate that the cardiolipin transacylase enzyme tafazzin plays a key role in regulating mouse B cell function and metabolic activity during activation.

## Introduction

Proper functioning of B lymphocytes is essential to establish a normal immune response [1]. Naïve B cells proliferate and differentiate upon activation in order to provide humoral responses and long term immunity. Lipopolysaccharide (LPS) is a gram-negative bacterial product that can activate murine B cells through binding to the Toll-like receptor-4 [2]. Activation of B cells with LPS induces B cell proliferation, chemokines synthesis and secretion and production of antibodies [3, 4]. Normal metabolic activity in B lymphocytes is critical for B cell homoeostasis and immune responses. Importantly, B lymphocytes increase oxygen consumption and lactate production as a response to activation and this is required to support proper B cell function. For example, activation of murine B cells with LPS has been shown to increase both mitochondrial oxidative phosphorylation and glycolysis [3].

Tafazzin (Taz) is a transacylase enzyme involved in remodeling and maturation of the mitochondrial phospholipid cardiolipin (CL) [5]. CL is an essential phospholipid of the inner mitochondrial membrane where it comprises approximately 15-20% of inner mitochondrial membrane phospholipid mass [6]. CL is required for activation of many enzymes of the electron transport chain [7]. Mutations in the tafazzin gene (TAFAZZIN) are observed in the rare (1:300,000) X-linked genetic disease Barth Syndrome (BTHS) [8, 9]. BTHS patient cells exhibit reduced ability to produce ATP from oxidative phosphorylation which leads to the dysfunction of metabolically active tissues such as the heart and skeletal muscle [10]. Although cardiomyopathy is the major clinical problem in BTHS, many of these patients additionally suffer from severe infections caused by neutropenia [8, 9]. The molecular mechanisms underlying this neutropenia remain unclear. Interestingly, neutrophils from BTHS patients exhibit normal motility, phagocytosis, mitochondrial shape and mass, with increased annexin-V binding in the absence of apoptosis [11]. It has also been observed that a few BTHS patients exhibit infections despite consistent prevention of neutropenia. Although B cells play a key role in the development of the immune response to infection, no studies have examined how Taz-deficiency impacts on B lymphocyte proliferation and function during their activation. B lymphocyte dysfunction may contribute to immune system dysfunction and potentially increase susceptibility to infection [12]. For example, it is well established that B cells play a critical role in neutrophil recruitment, activation of other immune cells including T cells, and long-term immune protection [4, 13]. In the current study, we determined how Taz-deficiency impacts on the function of LPS stimulated B cells. In addition, we identify novel protein targets involved in the Taz-mediated regulation of LPS stimulated B cell function.

## Material and Methods

### Experimental animals

All experimental procedures in this study were performed with approval of the University of Manitoba Animal Policy and Welfare Committee in accordance with the Canadian Council on Animal Care guidelines and regulations. This study is reported in accordance with ARRIVE guidelines. All animals were housed in pathogen-free facility (12 h light/dark cycle) at the University of Manitoba and had free access to food and water. Tafazzin knockdown (TazKD) mice were generated by mating transgenic male mice containing a doxycycline (dox) inducible tafazzin specific short-hair-pin RNA (B6.Cg-Gt(ROSA)26Sortm1(H1/tetO-RNAi:Taz,CAG-tetR)Bsf/ZkhuJwith female C57BL/6J mice from Jackson Laboratory (Boston, MA). Tafazzin knock down was maintained postnatally by administering dox (625 mg of dox/kg of chow) as part of the rodent chow (Catalog no. TD.01306) from Harlan (Indianapolis, USA) as described previously [14]. Male offspring were weaned at 3 weeks of age onto the above diet and wild type (WT) animals additionally received the dox containing diet. Male mice were used for the experiments between 10 and 14 weeks of age. Male mice positive for the tafazzin shRNA transgene were identified by PCR as described previously [14].

### Cell culture

Splenic naïve B lymphocytes were isolated by magnetic bead negative selection using EasySep™ Mouse B Cell Isolation Kit (Catalog no. 19854) from STEMCELL Technologies (Vancouver, BC). B cells were cultured in RPMI 1640 supplemented with 10% FBS, 1% antibiotic/antimycotic, and 2-mercaptoethanol at 37°C with 5% CO_2_. Subsequent to isolation B cells were stimulated or not with 10 µg/ml LPS (Catalog no. L2887) obtained from Sigma-Aldrich (Oakville, ON). Unless otherwise indicated, all other reagents used were of analytical grade and were obtained from either Thermo Fisher Scientific (Winnipeg, MB) or Sigma-Aldrich (Oakville, ON).

### Surface marker analysis

The expression of B cell surface markers cluster of differentiation 86 (CD86) and cluster of differentiation 69 (CD69) were measured using flow cytometry. B cells (2×10^6^) were cultured in 1 ml/well in 24-well plate in the presence 10 µg/ml LPS at 37°C in 5% humidified CO_2_ for 24 h. Subsequently, the B cells were stained with APC-conjugated anti-cluster of differentiation 19 (CD19) in combination with either PE-A labelled anti-CD69 or PE-CY7-A labelled anti-CD86. Anti-CD19, anti-CD69, anti-CD86 were obtained from BD Biosciences (Mississauga, ON). Cells were analyzed by flow cytometry using a BD FACSCanto II instrument. FlowJo software was used for data analysis.

### Analysis of proliferating and apoptotic cells

Proliferating and apoptotic cells were determined by cell cycle analysis using the Nicoletti assay [15]. B cells were treated with 10 µg/ml LPS for 48 h and cultured in 24-well plates. The cells were washed two times with PBS and suspended in 100 µl of ice cold PBS. Cells were then fixed in 5 ml of 70% ice-cold ethanol. After fixation the cells were stained with DNA fluorescent dye that quantitatively stains the DNA (40 µg/ml propidium iodide, 0.5 mg/ ml RNase A, 0.1% Triton X-100, 1% sodium citrate) for 1 h at 4°C in the dark. Then flow cytometry analysis was performed to detect red fluorescence (460 nm) for 10,000 cells using an Attune NxT Model AFC2 flow cytometer (Thermo Fisher Scientific). The number of proliferating cells was determined by counting the number of cells entering the S/G2 phase from the G0/G1 phase (non-proliferating phase). The percentage of apoptotic nuclei in sub-G1 section, healthy nuclei in G0/G1 section, and proliferating nuclei in S/G2 section were estimated by analysis of the DNA histogram and statistically compared between the samples [15].

### Measurement of chemokines and antibodies

Purified B cells were incubated in RPMI 1640 media supplemented with FBS, 2-mercaptoethanol, and 1% antibiotic/antimycotic. B cells were stimulated with 10 μg/ml LPS at 37°C with 5% CO_2_. After incubation for 24 h or 72 h, the supernatant fractions were collected to measure chemokines or antibodies concentrations, respectively. Immunoglobulin G (IgG), Immunoglobulin M (IgM), and the neutrophil attractant chemokines keratinocytes-derived chemokine (KC) and macrophage-inflammatory protein-2 (MIP-2) levels in cell culture supernatants were measured using isotype-specific enzyme-linked immunosorbent assay (ELISA) kits. IgG (total) mouse uncoated ELISA kit (Catalog no. 88-50400-22) and IgM mouse uncoated ELISA kit (Catalog no. 88-50470-22) were from Thermo Fisher Scientific (Winnipeg, MB). Mouse CXCL2/MIP-2 DuoSet ELISA (Catalog no. DY452-05) and mouse CXCL1/KC DuoSet ELISA (Catalog no. DY453-05) were obtained from R&D Systems (Minneapolis, MN).

### Proteasome and immunoproteasome activity assays

Immunoproteasome activity was determined by measuring the proteolytic activity of β1i/LMP2with S310 (Ac-PAL-AMC) as a fluorescent substrate using a commercial kit (Catalog no. 10008041) from Cayman Chemicals (Ann Arbor, MI). The fluorescent intensity was read at excitation = 345 nm and emission = 445 nm. To measure proteasome activity, the 20S subunit activity was determined using fluorescent substrate-SUC-LLVY-AMC (Catalog no. 10008041) from Cayman Chemicals (Ann Arbor, MI). The fluorescent intensity was read at excitation = 360 nm and emission = 480 nm.

### Metabolic assays

Oxygen consumption rate (OCR) and extracellular acidification rate (ECAR) were measured in B cells using a Seahorse XF24 Extracellular Flux Analyzer (Agilent Technologies) with kits from Agilent (Santa Clara, CA). Briefly, isolated WT and TazKD B cells were stimulated with LPS for 48 h. B cells were then seeded in XF24 24-well plates coated with Cell-Tak from BD Biosciences (Mississauga, ON) at a density of 4×10^5^ cells per well. The Cell-Tak (1.12 μg/well) was used to generate an adherent monolayer of B cells on the XF24 24-well plate. For mitochondrial function stress test (OCR assay) the following final concentrations of drugs were used: 1 μM oligomycin, 1 μM FCCP, 1 μM antimycin A, and 1 μM rotenone as per the manufacturer’s instructions. For glycolytic function test (ECAR assay) the following final concentrations of drugs were used: 10 μM glucose, 1 μM oligomycin, and 25 μM 2-DG as per the manufacturer’s instructions.

### Western blot analysis

Cell proteins were extracted using RIPA lysis buffer (Catalog no. 89900) from Thermo Fisher (Winnipeg, MB). The total protein levels were measured by the Bradford method [16]. Briefly, 30 µg of protein was mixed with an equal volume of 2×Laemmli buffer supplemented with 5% 2-mercaptoethanol (Catalog no. 1610737) then loaded in 10% polyacrylamide precast gel (TGX™ FastCast™ Acrylamide Starter Kit, 10%, Catalog no. 1610172) from BioRad Laboratories (Mississauga, ON). After 1 h electrophoresis, the proteins were transferred onto a polyvinylidene difluoride membrane at 4°C. This was followed by blocking the membrane using 5% milk in a solution of TBS containing 0.5% Tween 20 at room temperature for 1 h. Then, the membrane was washed with 0.5% Tween 20 three times and incubated with primary antibody overnight at 4°C. The primary antibody was removed the next day and the membrane was incubated with secondary antibody at room temperature for 1 h. Enhanced chemiluminescence substrate (Clarity Western ECL Substrate, Catalog no. 1705060) from Bio-Rad Laboratories (Mississauga, ON) was used to visualize the protein bands. The primary antibodies used for Western blotting were as follows: Tafazzin (Catalog no. sc-365810), PI 3-kinase p85α (Catalog no. sc-1637), Akt (Catalog no. sc-5298), p-Akt (Catalog no. sc-293125), and β-Actin (Catalog no. sc-47778) from Santa Cruz Biotechnolgies Inc. (Dallas, TX). Band intensity was quantified using Image J software.

### Proteomic analysis

Cell lysis protocol: B cells were resuspended in lysis buffer (100 mM HEPES, pH 8.5, 4% SDS, 1X protease inhibitors) and sonicated three times for 15 s per cycle with a 1 min cooling period (on ice) between each cycle. Then cell lysates were centrifuged at 15000g for 10 min to remove insoluble cellular debris.

Protein Digestion and Tandem Mass Tag (TMT) labeling: The Pierce detergent compatible Bradford assay kit from Thermo Fisher (Winnipeg, MB) was used to determine the protein concentration. All protein samples were processed and handled using single-pot solid-phase-enhanced sample preparation protocol. Prior to the sample preparation protocol treatment, two types of carboxylate-modified SeraMag Speed beads from GE Life Sciences (Mississauga, ON) were combined in a ratio of 1:1 (v/v), rinsed, and reconstituted in water at a concentration of 20 μg solids/μL. Initially, 100 µg of lysate was reduced with 10 mM (final concentration) DTT for 30 minutes at 60°C followed by alkylation using 50 mM (final concentration) IAA for 45 min in dark at room temperature. After that, 10 μL of the prepared bead mix was added to the lysate and samples were adjusted to pH 7.0 using HEPES buffer. Acetonitrile was added to a final concentration of 70% (v/v) to promote proteins binding to the beads. Samples were then incubated at room temperature on a tube rotator for 18 min. Subsequently, beads were immobilized on a magnetic rack for 1 min. The pellet was rinsed 2X with 200 µl of 70% ethanol and 1X with 200 µl of 100% acetonitrile while on the magnetic rack and the supernatant was discarded. Rinsed beads were resuspended in 40 μL of 50 mM HEPES buffer, pH 8.0, supplemented with trypsin at an enzyme-to-protein ratio of 1:25 (w/w) and incubated for 16 h at 37°C. After overnight digestion, peptides were eluted and immediately labeled with TMT tags. After overnight digestion, peptides were eluted and immediately labeled with TMT tags. TMT labeling was performed as specified by the manufacturer’s instructions (Thermo Fisher, Winnipeg, MB), except that TMT tags were dissolved in DMSO. Equivalent labeled samples within each TMT set were mixed prior to 1D LC/MS/MS.

Mass spectrometry data acquisition: Analysis of TMT labeled peptide digests was carried out on an Orbitrap Q Exactive HF-X instrument from Thermo Fisher Scientific (Bremen, DE). The sample was introduced using an Easy-nLC 1000 system from Thermo Fisher Scientific (Bremen, DE) at 2 μg per injection. Data acquisition on the Orbitrap Q Exactive HF-X instrument was configured for data-dependent method using the full MS/DD-MS/MS setup in a positive mode.

### Statistical analysis

All experimental results were expressed as mean ± SD. The levels of significance were calculated using two tailed unpaired Student’s t test unless otherwise specified one-way-analysis of variance followed by multiple comparisons using the Tukey’s test. A *p* value of <0.05 was considered statistically significant.

## Results

### TAZ-deficiency decreases proliferation and reduces survival markers in LPS stimulated B cells

The percentage of cells in the Sub-G1, G0/G1 and S/G2 phases of the cell cycle were determined in naïve WT B cells and TazKD B cells and in these cells incubated with LPS for 48 h. The number of cells in Sub-G1, G0/G1 and S/G2 was unaltered in naïve TazKD B cells compared to naïve WT B cells (Fig. 1A). In contrast, LPS stimulation of TazKD B cells resulted in an increase of cells in G0/G1 (*p*=0.0142) and a decrease in S/G2 (*p*=0.0016) compared to WT B cells (Fig. 1B). We confirmed that Taz protein was significantly reduced approximately 50% (*p*<0.001) in naïve B cells isolated from TazKD animals and that Taz protein remained reduced in LPS stimulated B cells (*p*<0.0001) (Fig. 1C). Activation of phosphoinositide 3-kinase (PI3K) is essential for B cell function and proliferation. Toll-like receptor-4 agonists such as LPS trigger PI3K activation and promote activation of the AKt signaling pathway [17]. PI3K, AKt, and PAKt protein expression were examined in naïve and LPS stimulated WT and TazKD B cells. PI3K protein expression was decreased 40% (*p*=0.0229) in naïve TazKD B cells and 70% (*p*<0.0001) in LPS stimulated TazKD B cells compared to WT B cells, respectively. A greater reduction in PI3K protein expression was observed in LPS stimulated TazKD B cells compared to naïve TazKD B cells (*p*=0.0248). Basal AKt protein levels were unaltered in naïve and LPS stimulated TazKD B cells compared to naïve and LPS stimulated WT B cells, respectively. In contrast, pAKt protein levels were reduced 35% (*p*=0.0246) in naïve TazKD B cells and 80% (*p*<0.0001) in LPS stimulated TazKD B cells compared to naïve and LPS stimulated WT B cells, respectively. Although the ratio of pAKt/AKt protein was reduced minimally in naïve TazKD B cells compared to naïve WT B cells (*p*=0.0735), LPS stimulation resulted in a 65% reduction (*p*=0.0002) in the ratio of pAKt/AKt protein in TazKD B cells compared to WT B cells. These data suggest that Taz-deficiency reduces proliferation and PI3K and AKt pathway activation in LPS activated B cells.

**Figure 1.**
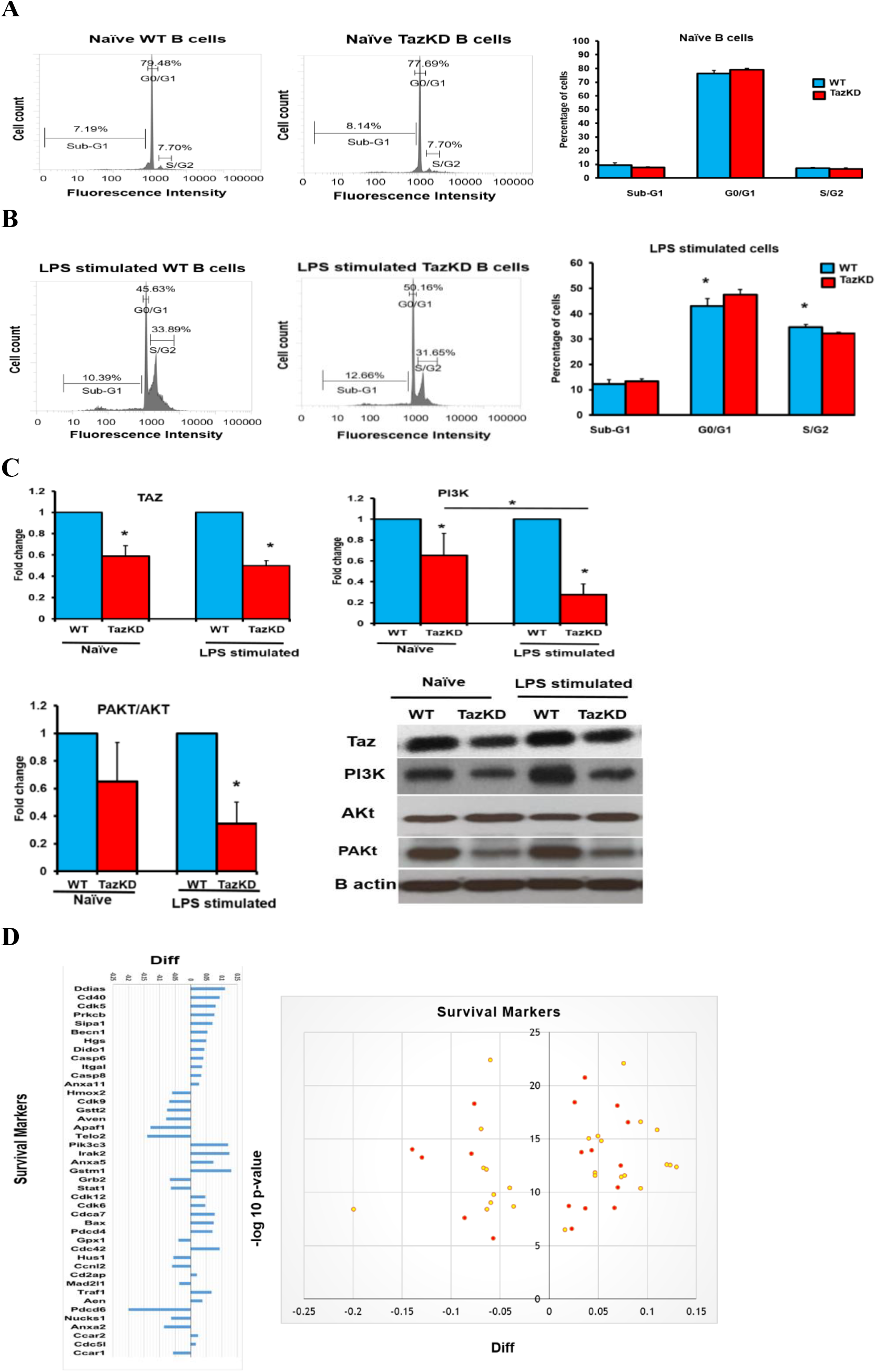
Taz-deficiency decreases cell proliferation and reduces cell survival markers in LPS stimulated B cells. The percentage of cells in Sub-G1, G0/G1 and S/G2 phases of the cell cycle and the protein levels of Taz, PI3K, Akt, and PAKt were determined in naïve WT and TazKD B cells or WT and TazKD B cells stimulated for 48 h with LPS as described in Materials and Methods. **A**. Flow cytometry representative figures of naïve WT and TazKD B cells and percentage of cells in Sub-G1, G0/G1 and S/G2, n=4-6. **B**. Flow cytometry representative figures of LPS stimulated WT and TazKD B cells and percentage of cells in Sub-G1, G0/G1 and S/G2, n=4-6. \**p*<0.05 compared with WT B cells. **C**. Protein levels of Taz, PI3K, Akt, and pAKt and B actin as loading control, n=4. \**p*<0.05 compared with WT B cells. **D**. Survival markers in LPS stimulated WT and TazKD B cells. Volcano plot and log2 fold change ratios of survival protein markers in LPS stimulated WT and TazKD B cells are shown. Significantly changed proteins are indicated in red. n=3, *p*<0.05 compared with WT B cells.

Proteomic analysis indicated alteration in the levels of several survival and apoptosis markers in LPS stimulated TazKD B cells compared to LPS stimulated WT B cells (Fig. 1D). This included upregulation in the apoptotic markers Death-Inducer Obliterator 1, Beclin1, apoptosis regulator BAX, Caspase-6, Caspase-8, and Annexin A11. In addition, Aven, an anti-apoptotic protein that binds BCL-XL and impairs the activation of many Caspases [18], was down regulated in LPS stimulated TazKD B cells compared to WT B cells. In addition, the key cell cycle regulator cyclin-dependent kinase 9 [19] was down regulated in LPS stimulated TazKD B cells compared to WT B cells. Cyclin-dependent kinase 9 is serine/threonine kinase required for the formation of the catalytic core of the transcription elongation factor b and regulates the S phase of the cell cycle, proliferation, and differentiation of many cell types [20]. Transcription elongation factor b recruits RNA polymerase II and thus stimulates the transcription and elongation of many genes. Similarly, the level of the cell cycle checkpoint protein HUS1,which negatively affects cell cycle control [21], was reduced in LPS stimulated TazKD B cells compared to WT B cells as was telomere length regulation protein 2. Telomere length regulation protein 2 is an essential protein for telomere length maintenance and thus prevents cellular senescence and favors the proliferation and survival of many cells [22]. Telomere length regulation protein 2 enhances the stability of phosphatidylinositol 3-kinase-related protein kinases. In addition, signal transducer and activator of transcription 1 was decreased in LPS stimulated TazKD B cells compared to WT cells. Signal transducer and activator of transcription 1 is a transcription factor necessary for the differentiation of naïve B cells into IgM antibody-producing marginal zone B cells thus ensuring a quick response to infectious pathogens and is essential for the transcription of many genes involved in immune cell division, survival, activation and recruitment [23]. These data suggest that Taz-deficiency impacts the survival and proliferation of LPS activated B lymphocytes.

### TAZ-deficiency reduces the expression of surface markers and immunogenic markers in LPS stimulated B cells

The above data indicated that Taz-deficiency impacts B cell proliferation. B lymphocytes are antigen-presenting cells and the ability of B cells to upregulate surface markers expression is essential for intercellular communication. Expression of the costimulatory molecule CD86 in B cells is required to initiate interaction with T lymphocytes [24]. In addition, activated B cells exhibit CD69 induction, a marker of early leukocyte activation [25]. We measured CD86 and CD69 expression on the surface of naïve and LPS stimulated WT and TazKD B cells. As expected, minimal expression of CD86 and CD69 was observed in both naïve WT and TazKD B cells and Taz deficiency did not impact their expression (Figs. 2A, 2B). In contrast, CD86 and CD69 expression were markedly upregulated in both WT B cells and TazKD B cells stimulated with LPS (Figs. 2C, 2D). However, we observed a 10% reduction (*p*=0.0172) in CD86 expression and an 15% reduction (*p*=0.0027) in CD69 expression in LPS stimulated TazKD B cells compared to LPS stimulated WT B cells (Figs. 2C, 2D).

**Figure 2.**
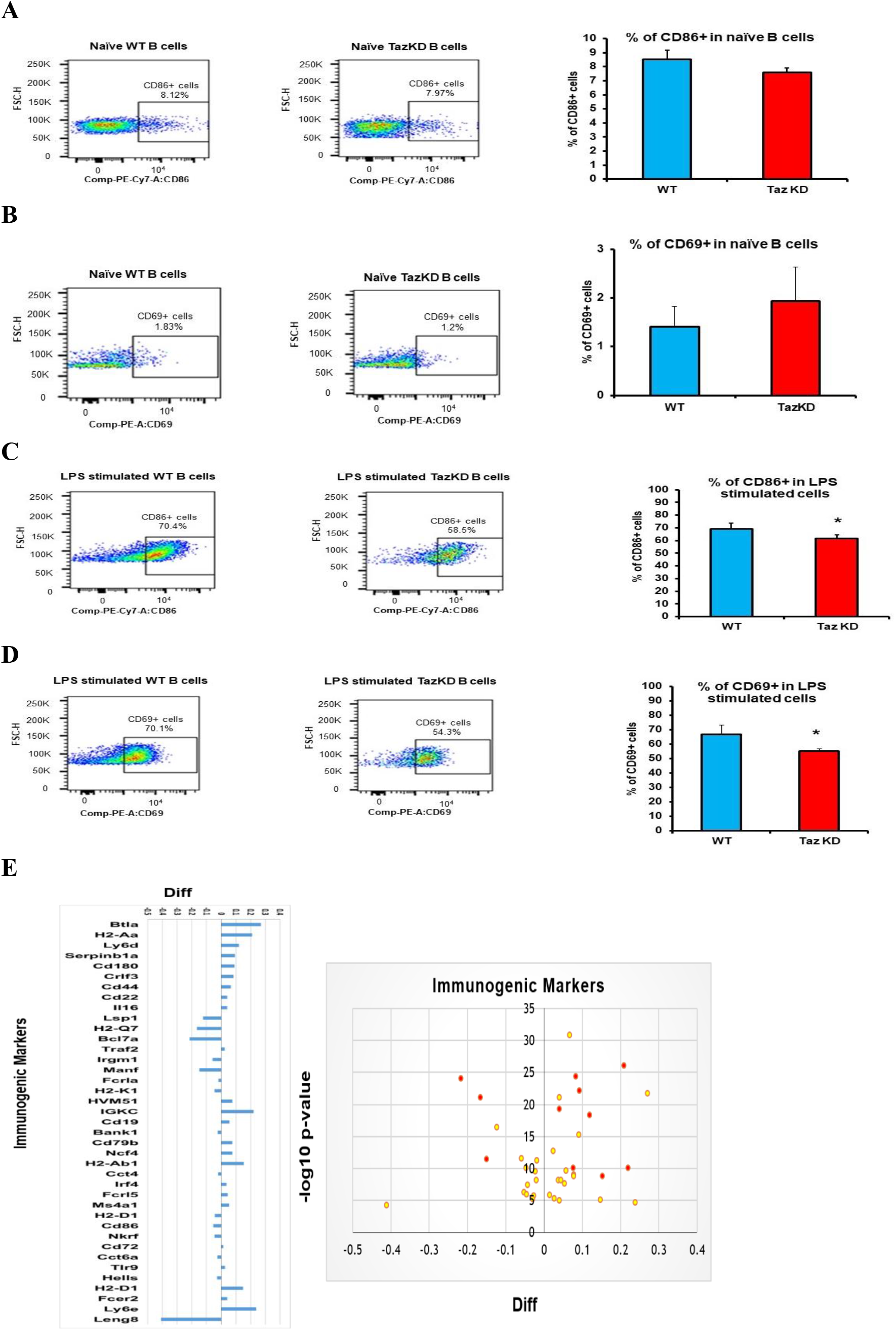
Taz-deficiency reduces the expression of B cell surface markers and immunogenic markers. The percent of CD86+ and CD69+ expressing cells was determined in naïve WT and TazKD B cells or WT and TazKD B cells stimulated for 48 h with LPS as described in Materials and Methods. Representative flow cytometry histograms are shown in A-D. **A**. CD86+ in naïve WT and TazKD B cells. **B**. CD69 expression in naïve WT and TazKD B cells. **C**. CD86+ expression in LPS stimulated WT and TazKD B cells. **D**. CD69+ expression in LPS stimulated WT and TazKD B cells. n=3, \**p*<0.05 compared with WT B cells. **E**. Immunogenic markers in LPS stimulated WT and TazKD B cells. Volcano plot and log2 fold change ratios of immunogenic markers in LPS stimulated WT and TazKD B cells is shown. Significantly changed proteins are indicated in red. n=3, *p*<0.05 compared with WT B cells.

Proteomic analysis indicated alteration in the level of several other B lymphocyte related receptors and markers in stimulated TazKD B cells compared to WT B cells (Fig. 2E). For example, lymphocyte-specific protein 1, a cytoskeletal protein expressed in B lymphocytes and other immune cells required for the migration of these cells to the site of inflammation [26], was reduced in LPS stimulated TazKD B cells compared to WT B cells. Immunity-related GTPase family M protein 1, a member of a family of interferon-inducible cytoplasmic immunity-related GTPases required for mitochondrial homeostasis in B lymphocytes [27], was reduced in LPS stimulated TazKD B cells compared to WT B cells. Deficiency in immunity-related GTPase family M protein 1 is known to weaken the adaptive immune system and phagosome maturation and was reduced in LPS stimulated TazKD B cells compared to WT B cells. In contrast, the level of B- and T-lymphocyte attenuator was increased in LPS stimulated TazKD B cells compared to WT B cells. B- and T-lymphocyte attenuator is an “off” signal that inhibits the maturation of B-lymphocytes and is expressed in high amounts in naïve and immature B-lymphocytes in comparison with active and mature B-lymphocytes [28]. In addition, the level of cluster of differentiation-22 was increased in LPS stimulated TazKD B cells compared to WT B cells. Cluster of differentiation-22 confines B cell differentiation to plasmablasts, inhibits interleukin-6 production and increases interleukin-10 production, thus inducing an immune-tolerance state [29, 30]. Finally, pro-interleukin-16 was increased in LPS stimulated TazKD B cells compared to WT B cells. Pro-interleukin-16 is a mediator of apoptosis through activation of caspase-3 in B lymphocytes [31]. These data suggest that Taz-deficiency alters expression of B lymphocyte receptors and immunogenic markers.

### TAZ-deficiency reduces secretion of chemokines and IgM in LPS stimulated B cells

Activated B cells synthesize and secret antibodies. Cells from WT and TazKD mice were isolated and stimulated with LPS for 72 h the supernatants collected and level of the secreted antibodies IgG and IgM determined. The level of IgG in the supernatant was unaltered in LPS stimulated TazKD B cells compared to WT B cells (Fig. 3A). In contrast, the level of IgM was reduced 20% (*p*<0.0001) in the supernatant of LPS stimulated TazKD B cells compared to WT B cells (Fig. 3B). Activated B cells also secrete chemokines that promote leukocyte recruitment to sites of infections in order to increase the immune response. Murine B cells synthesize and secrete two CXC chemokines, cytokine-induced neutrophil chemoattractant (KC) and macrophage inflammatory protein-2 (MIP-2) in response to LPS [4]. B cells from WT and TazKD mice were isolated and stimulated with LPS for 24 h the supernatants collected and level of secreted KC and MIP-2 determined. The level of KC was decreased 55% (*p*<0.0001) and the level of MIP-2 was decreased 50% (*p*<0.0001) in the supernatant of LPS stimulated TazKD B cells compared to WT B cells (Fig. 3C, 3D). Thus, Taz-deficiency reduces secretion of KC and MIP-2 chemokines and IgM in LPS stimulated B cells.

**Figure 3.**
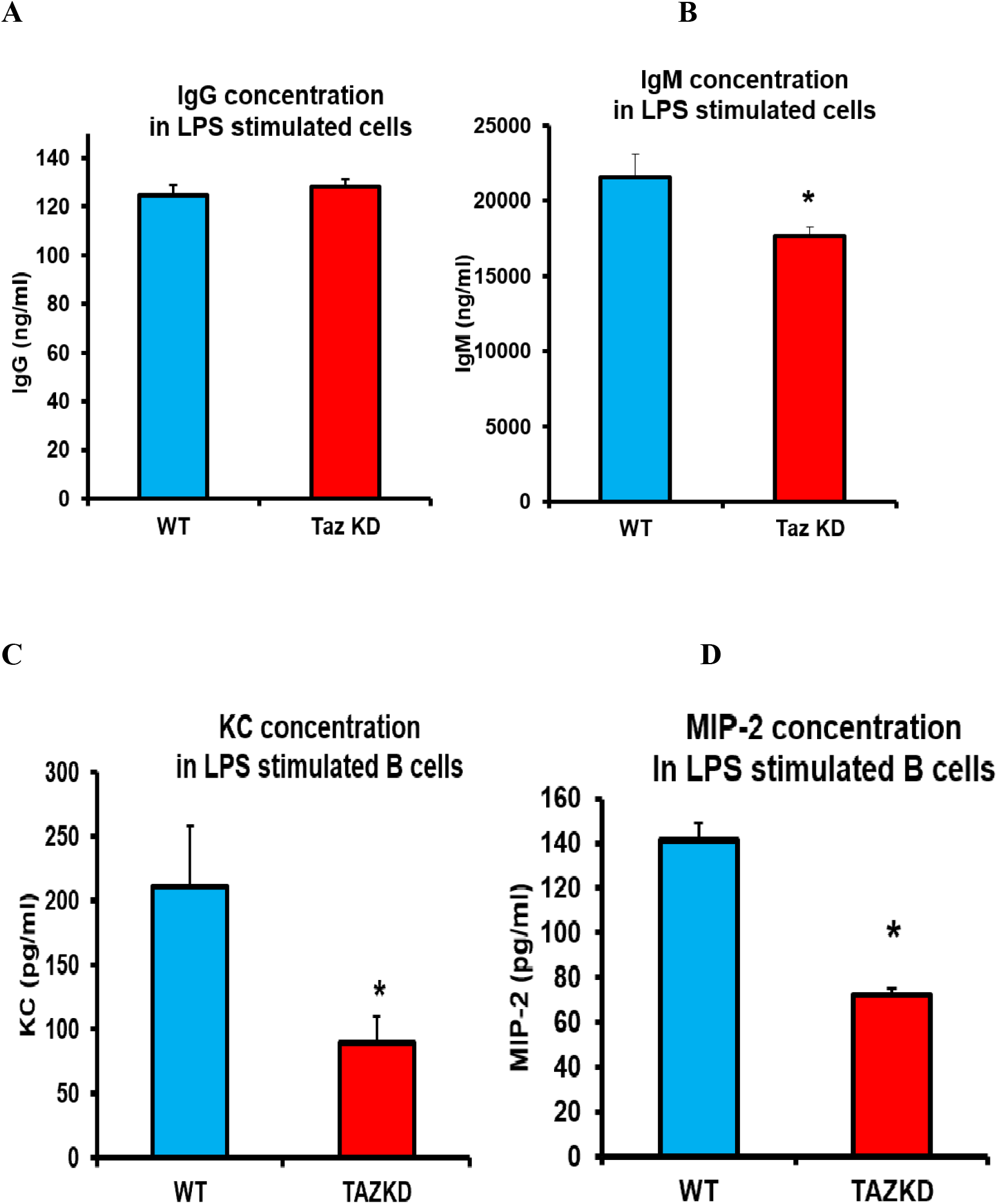
Taz-deficiency reduces secretion of chemokines and IgM in LPS stimulated B cells. B cells were isolated from WT or TazKD mice and stimulated with LPS for 72 h for measurement of antibodies concentrations or stimulated for 24 h with LPS for measurement of chemokines concentrations as described in Materials and Methods. **A**. IgG. **B**. IgM. **C**. KC. **D**. MIP-2. n=4-6, \**p*<0.05 compared with WT B cells.

### TAZ-deficiency reduces proteasome and immunoproteasome activities in LPS stimulated B cells

The ubiquitin proteasome system is the main cellular protein degradation pathway which degrades damaged proteins by proteolysis and it is an ATP-consuming process [32]. Inactivation of the 26S proteasome inhibits activation and function of many immune cells including B lymphocytes [33, 34]. Proteasome activity was determined in naïve B cells from WT and TazKD mice or WT and TazKD B cells stimulated with LPS for 24 h. Proteasome activity was reduced approximately 15% (*p*<0.011) in naïve TazKD B cells compared to naïve WT B cells (Fig. 4A). In addition, proteasome activity was decreased approximately 60% (*p*<0.0001) in LPS stimulated TazKD B cells compared to LPS stimulated WT B cells (Fig. 4B). The reduction in proteasome activity was greater in LPS stimulated TazKD B cells compared to naïve TazKD B cells (Fig. 4A, B).

**Figure 4.**
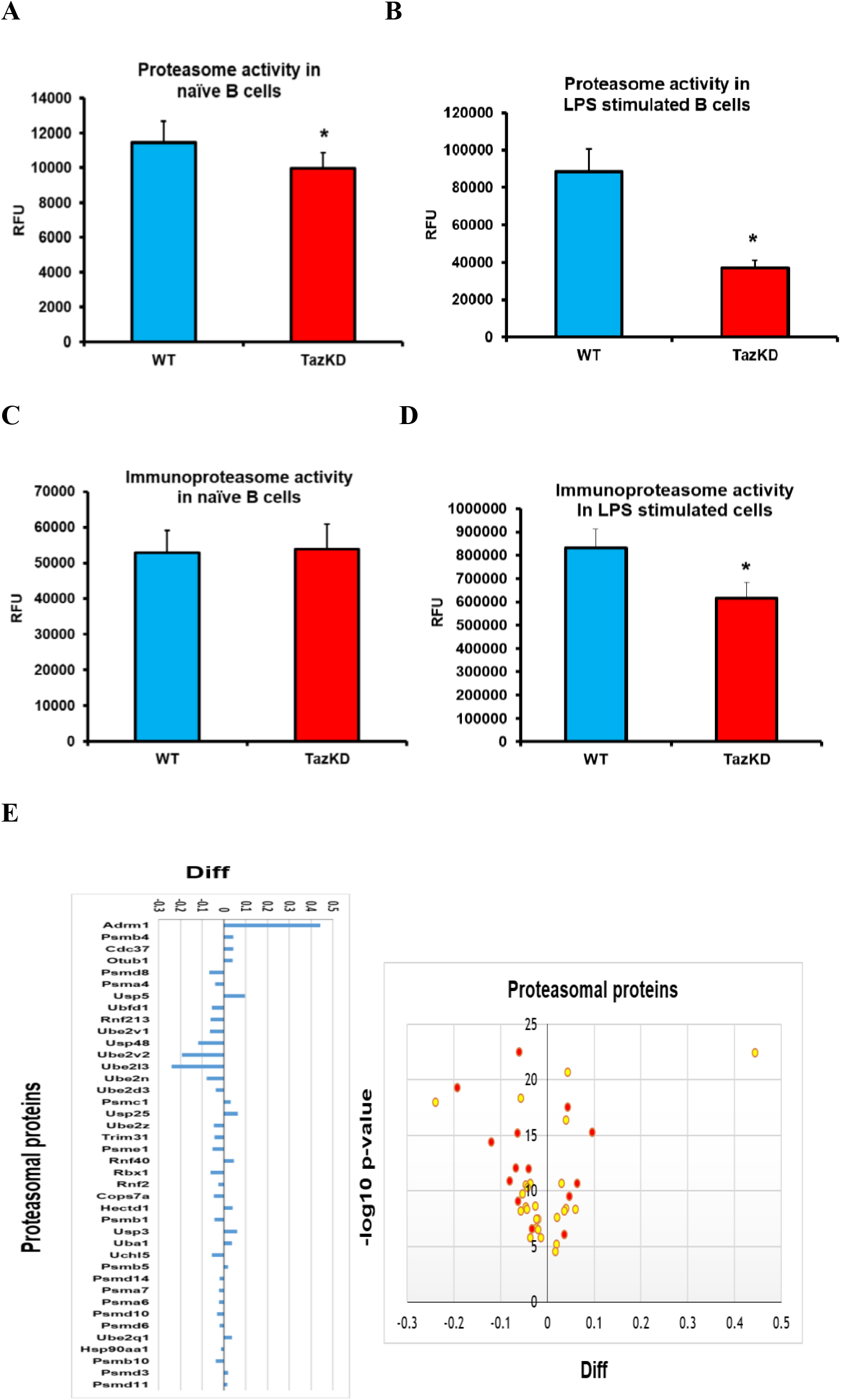
TAZ-deficiency reduces proteasome and immunoproteasome activity in LPS stimulated B cells. Proteasome and immunoproteasome activities were determined in naïve B cells from WT and TazKD mice or WT and TazKD B cells stimulated with LPS for 24 h as described in Materials and Methods. **A**. Proteasome activity in naïve WT and TazKD B cells. **B**. Proteasome activity in LPS stimulated WT and TazKD B cells. **C**. Immunoproteasome activity in naïve WT and TazKD B cells. **D**. Immunoproteasome activity in LPS stimulated WT and TazKD B cells. n=4, \**p*<0.05 compared with WT B cells. **E**. Proteasomal proteins in LPS stimulated WT and TazKD B cells. Volcano plot and log2 fold change ratios of proteasomal proteins in LPS stimulated WT and TazKD B cells are shown. Significantly changed proteins are indicated in red. n=3, *p*<0.05 compared with WT B cells.

The immunoproteasome is a specialized form of proteasome that is crucial for antigen presentation performed by various activated immune cells [35]. Immunoproteasome activity was determined in naïve B cells from WT and TazKD mice or WT and TazKD B cells stimulated with LPS for 24 h. Immunoproteasome activity was unaltered in naïve TazKD B cells compared to naïve WT B cells (Fig. 4C). In contrast, immunoproteasome activity was reduced 25% (*p*<0.0001) in LPS stimulated TazKD B cells compared to LPS stimulated WT B cells (Fig. 4D).

Proteomic analysis indicated alteration in the level of several B lymphocyte proteasomal proteins in LPS stimulated TazKD B cells compared to LPS stimulated WT B cells (Fig. 4E). in These included a reduction in proteasome activator complex subunit 1 and the induced proteolytic core subunit proteasome subunit beta type-10 which are linked to reduced function of the immunoproteasome and thus a defect in antigen presentation [36]. Ubiquitin carboxyl-terminal hydrolase 48 promotes survival of cells by stabilizing their genetic content [37] and was reduced in LPS stimulated TazKD B cells compared to WT B cells. The 26S proteasome recognizes misfolded proteins or proteins assigned for degradation by tagging these proteins with ubiquitin proteins [38]. This ubiquitination requires the combined function of ubiquitin activating enzymes E1, ubiquitin conjugating enzymes E2, and E3 ligases [39]. Several ubiquitination enzymes exhibited reduced level of expression in LPS stimulated TazKD B cells compared to LPS stimulated WT B cells. These included the conjugating enzymes ubiquitin-conjugating enzyme E2 variant-1, -2, and ubiquitin-conjugating enzyme E2 L3, and the E3 ligase ring finger protein 213. In addition, many 19S proteasome subunits, deubiquitinases and ubiquitin receptors required for binding ubiquitinated tagged proteins [40] were decreased in LPS stimulated TazKD B-Lymphocytes compared to LPS simulated WT B cells including 26S proteasome non-ATPase regulatory subunit 8, 26S proteasome non-ATPase regulatory subunit 14, and 26S proteasome non-ATPase regulatory subunit 10. These observed decreases in ubiquitin-26S proteasome related proteins coincided with the reduced proteasomal and immunioproteosomal activities in LPS stimulated TazKD B cells. Thus, Taz-deficiency reduces proteasomal and immunoproteasome activities in LPS stimulated B cells.

### TAZ-deficiency reduces metabolic activity in LPS stimulated B cells

Naïve B cells are metabolically quiescent whereas LPS activation of B cells initiates a metabolic reprogramming which includes the augmentation of mitochondrial oxidative phosphorylation (OXPHOS) required for B cell proliferation and function [3]. Oxygen consumption rates (OCR) were determined in WT B cells and TazKD B cells after stimulation with LPS for 48 h using a Seahorse XF24 analyzer. Basal respiration (measured as OCR before adding inhibitors) was reduced 25% (*p*<0.0001) in LPS stimulated TazKD B cells compared to WT B cells (Fig. 5A). Maximal respiration (measured as OCR after adding FCCP an uncoupler of the electron transport chain) was approximately 25% lower (*p*<0.0001) in LPS stimulated TazKD B cells compared to WT B cells. In addition, spare respiratory capacity (calculated by the difference between the maximal OCR and the basal OCR) was reduced 35% (*p*<0.0001) in LPS stimulated TazKD B cells compared to WT B cells.

**Figure 5.**
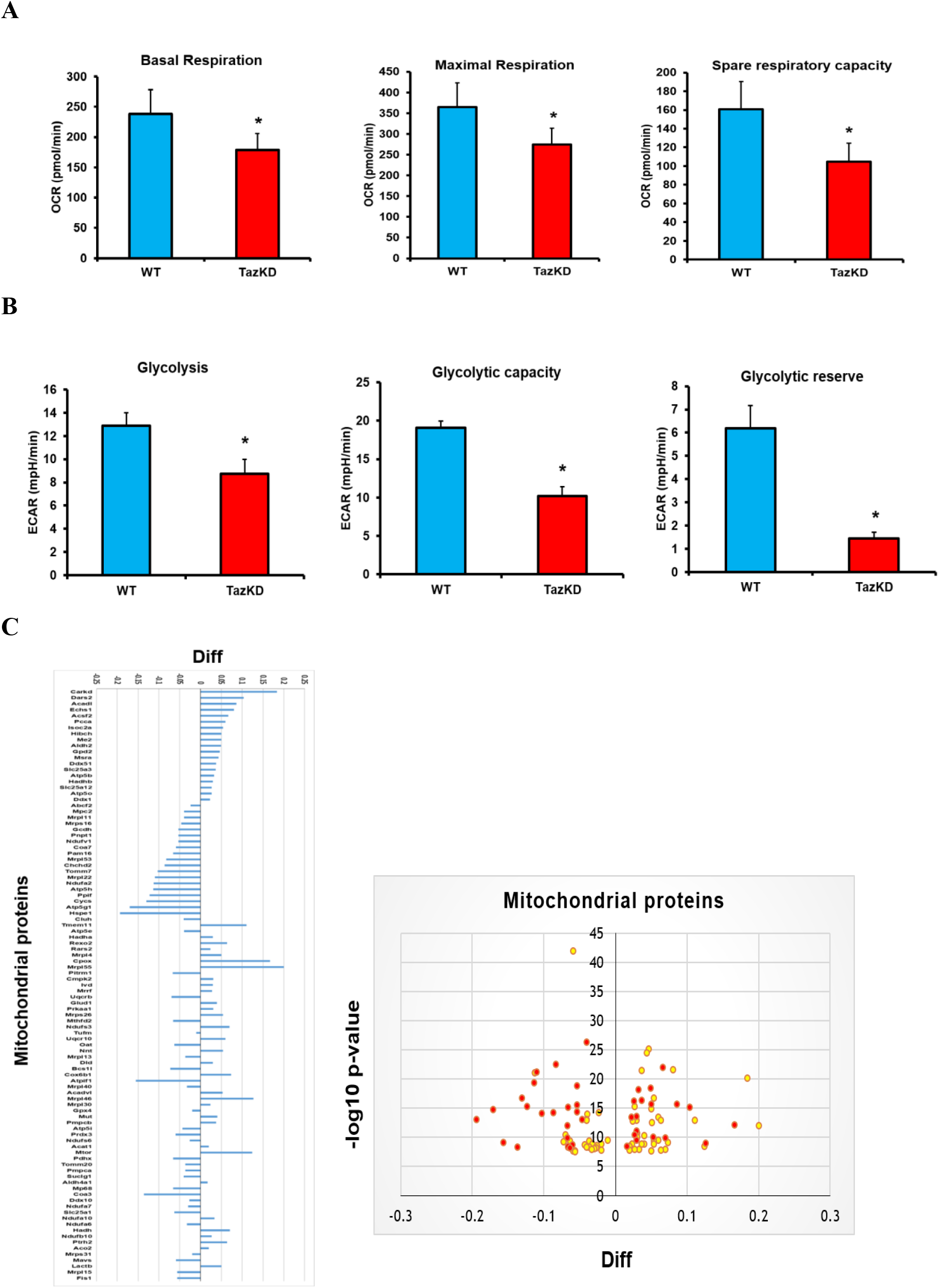
Taz-deficiency reduces metabolic activity in LPS stimulated B cells. Mitochondrial function and glycolytic activity were analyzed in LPS stimulated WT and TazKD B cells using a Seahorse XF24 analyzer as described in Materials and Methods. **A**. Basal respiration, maximal respiration and spare respiratory capacity. **B**. Glycolysis, glycolytic capacity and glycolytic reserve. n=3, \**p*<0.05 compared with WT B cells. **C**. Mitochondrial proteins in LPS stimulated WT and TazKD B cells. Volcano plot and log2 fold change ratios of mitochondrial proteins in LPS stimulated WT and TazKD B cells are shown. Significantly changed proteins are indicated in red n=3, *p*< 0.05 compared with WT B cells.

Previous studies have demonstrated that naïve B cells increase glycolysis upon LPS stimulation [41]. Glycolytic activity was determined in WT B cells and TazKD B cells after stimulation with LPS for 48 h using a Seahorse XF24 analyzer. Glycolysis (measured by determination of the change in extracellular acidification rate (ECAR) before oligomycin addition minus ECAR before glucose addition) was reduced 30% (*p*<0.0001) in LPS stimulated TazKD B cells compared to WT B cells (Fig. 5B). Glycolytic capacity (calculated as the difference between ECAR following addition of oligomycin minus basal ECAR), was reduced 50% (*p*<0.0001) in LPS stimulated TazKD B cells compared to WT B cells. Finally, glycolytic reserve (calculated as the difference between glycolytic capacity and glycolysis) was decreased 75% (*p*<0.0001) in LPS stimulated TazKD B cells compared to WT B cells (Fig. 5B). Thus, TAZ-deficiency reduces metabolic activity in LPS stimulated B cells.

Proteomic analysis indicated significant alteration in the level of many B lymphocyte mitochondrial proteins in LPS stimulated TazKD B cells compared to WT B cells (Fig. 5C). Among these included reduction in NADH dehydrogenase (ubiquinone) flavoprotein 1& 2 core subunits in mitochondrial Complex 1, the first of five mitochondrial complexes involved in the oxidative phosphorylation cascade [42]. Cytochrome c oxidase assembly factor 7 is one of the 14 assembly subunits required for the formation of functional cytochrome c oxidase Complex IV [43]. Cytochrome c oxidase assembly factor 7 levels were lower in LPS stimulated TazKD B cells compared to WT B cells. Translocase of the outer membrane 7, an essential assembly subunit for the translocase of the outer membrane complexes and translocase of the outer membrane 20 and translocase of the outer membrane 22 [44], were reduced in LPS stimulated TazKD B cells compared to WT B cells. The level of mitochondrial import inner membrane translocase subunit TIM16, which forms a complex with mitochondrial import inner membrane translocase subunit TIM14 to facilitate its insertion via hydrophobic N-terminal domain with the presequence translocases of the inner mitochondrial membrane [45], was reduced in LPS stimulated TazKD B cells compared to WT B cells. Finally, the level of cytochrome c, which is a main participant in the electron transport chain of mitochondria, and ATP synthase required for ATP production in oxidative phosphorylation (ATP synthase F1 subunit beta), were reduced in LPS stimulated TazKD B cells compared to WT B cells. Thus, Taz-deficiency reduces mitochondrial function in LPS stimulated TazKD B lymphocytes.

## Discussion

B lymphocytes are responsible for humoral immunity and play a key role in the immune response and optimal mitochondrial function is required to support activated B cell function [2]. In this study, we examined how deficiency of Taz, a cardiolipin remodeling enzyme required for mitochondrial function, alters the metabolic activity of B cells and their response to activation by LPS in mice. We have demonstrated for the first time that Taz-deficiency in LPS activated mouse B cells results in impaired metabolic function and this may contribute to reduced B cell function.

Mitochondria play a vital role in the activation, survival, development, and differentiation of B lymphocytes and their response to various phenotypic and environmental changes [46]. Mitochondria are a highly dynamic organelle that relies on many proteins and enzymes to maintain their membrane composition, membrane potential, and control continuous rounds of fusion and fission in order to remain fully competent for respiration and ATP synthesis [47]. Naïve B cells predominately exist in a quiescent state until stimulation [46]. Activated B cells are metabolically reprogrammed resulting in an increase in both glycolysis and OXPHOS pathways required for rapid proliferation and differentiation [3]. It is well established that TAFAZZIN mutation is associated with mitochondrial dysfunction in different organs of BTHS patients including the heart and skeletal muscle [10]. Moreover, multiple studies have reported that Taz-deficiency in TazKD mice result in decreased mitochondrial respiration in several tissues [48, 49]. We previously reported that Epstein-Barr virus transformed human BTHS lymphoblasts exhibited impairment in mitochondrial function [50]. Herein, we demonstrate for the first time that LPS stimulated B cells isolated from TazKD mice exhibit a marked impairment in OXPHOS. In addition, our proteomic analysis identified a decrease in several mitochondrial proteins including NADH dehydrogenase (ubiquinone) flavoprotein 1& 2 core subunits in mitochondrial Complex 1, cytochrome c oxidase assembly factor 7, translocase of the outer membrane 7, import inner membrane translocase subunit TIM16, cytochrome c, and ATP synthase in LPS stimulated TazKD B cells compared to LPS stimulated WT B cells. Collectively, a reduction in the above mitochondrial proteins and impairment in OXPHOS indicate a marked defect in mitochondrial function of LPS activated B cells from TazKD mice.

Activation of B cells with LPS increases cell proliferation, induces surface marker expression, chemokine signaling, and promotes B cell differentiation into antibody secreting cells required to amplify the immune response [3,4]. PI3K activation is essential for B cell proliferation and is initiated upon B cell stimulation. The AKt serine/threonine kinase is one of the critical signaling molecules of the PI3K activation pathway. Increasing evidence suggests that reduced activity of the PI3K pathway and insufficient AKt activation inhibits glucose utilization in B cells [17]. For example, inhibition of PI3K activity in WT B cells by LY294002 treatment was shown to diminish glycolysis [51]. Consistent with attenuation of PI3K activity in LPS stimulated TazKD B cells, we observed impaired glycolytic activity. In B cells, glycolysis is a crucial for their proliferation and antibody production and inhibiting glycolysis leads to impaired B cell growth and suppresses antibody secretion [3]. Our observation of a reduced number of cells entering the S/G2 phase of the cell cycle and reduced IgM secretion in LPS stimulated TazKD B cells is consistent with these results. In addition, we observed a reduction in CD69 and CD86 surface marker expression in LPS stimulated TazKD B cells which may potentially result in reduced intercellular communication with other cells of the immune system. For example, MIP-2 and KC, CXC family member chemokines, are secreted by activated murine B cells and are known to recruit neutrophils to infection sites [4]. B cells isolated from TazKD mice exhibited a reduced ability to produce MIP-2 and KC after LPS activation. Thus, decreased MIP-2 and CK secretion by TazKD B cells could potentially inhibit neutrophil recruitment and increase susceptibility to infection. Together, these results indicate that LPS activated B cells from TazKD mice exhibit a reduced function in their ability to amplify the immune response.

The ubiquitin proteasome system is the main protein degradation pathway which degrades damaged proteins by proteolysis [35]. The 26S proteasome is a multiprotein complex and is considered the master protein degradation machinery in cells [52]. The 26S proteasome degradation machinery regulates many cellular processes including proliferation, DNA repair, cell cycle, antigen presentation, and autophagy. The 26S proteasome consists of two large complexes, the 19S proteasome or the regulatory complex and the 20S proteasome where the proteolytic core is localized. Intact function of the 26S proteasome requires strong binding between the 19S proteasome and 20S proteasome and this assembly is crucial for optimal degradation function of the 26S proteasome complex [53]. Many autoimmune diseases, neurodegenerative disorders and cancers can be linked to alteration in 26S proteasome assembly and function. It was previously reported that inactivation of the 26S proteasome inhibits the function of many immune cells including B lymphocytes resulting in their apoptosis [34]. Moreover, mitochondrial dysfunction is induced by defective 26S proteasome machinery [55]. In addition, efficient and intact 26S proteasome activity is required to remove damaged proteins from the mitochondria and maintain mitochondrial homeostasis [55,56]. The immunoproteasome is a specialized form of proteasome crucial for the antigen presentation elicited by different immune cells [54]. It is constitutively expressed in immune cells especially in antigen-presenting cells such as B lymphocytes. The immunoproteasome is composed of a cap known as the proteasome activator complex subunit 1 or 11S proteasome and an induced form of the 20S proteasome that exhibits a specialized proteolytic core for antigenic proteins [52]. It was previously demonstrated that immunoproteasome inhibition impairs naïve B cell activation [35]. Consistent with reduced proteasome and immunoproteasome activities, our proteomic data revealed for the first time that LPS stimulated TazKD B cells exhibited a significant reduction in the level of 26S proteasome subunits and chaperones. The observed alteration in proteasome subunits might contribute to the mitochondrial dysfunction in LPS stimulated TazKD B cells. In addition, several 19S proteasome subunits, deubiquitinases and ubiquitin receptors required for binding the ubiquitinated tagged proteins were decreased in LPS stimulated TazKD B-Lymphocytes. A decrease in these proteasome related proteins in LPS stimulated TazKD B cells may additionally contribute to reduced activity of the proteasome machinery and survival of these cells.

In conclusion, we have demonstrated for the first time that LPS activated TazKD B cells exhibit reduced mitochondrial OXPHOS and glycolysis, reduced proliferation, lowered expression of CD86 and CD69 surface markers, reduced secretion of IgM antibody and the KC and MIP-2 chemokines, and reduced proteasome and immunoproteasome activities. In addition, our proteomic analysis revealed striking alterations in protein targets that regulate cell survival, immunogenicity, proteasomal processing and mitochondrial function consistent with these functional observations. These studies highlight a novel role for Taz in the maintenance of normal B cell function.

## List of abbreviations

Taz: tafazzin
LPS: lipopolysaccharide
CL: cardiolipin
BTHS: Barth Syndrome
WT: wild type
TazKD: tafazzin knockdown
CD86: cluster of differentiation 86
CD69: cluster of differentiation 69
KC: keratinocytes-derived chemokine
MIP-2: macrophage-inflammatory protein-2
IgG: Immunoglobulin G
IgM: Immunoglobulin M
OCR: oxygen consumption rate
ECAR: extracellular acidification rate
TMT: tandem mass tag
PI3K: phosphatidylinositol-3-kinase
AKt: protein kinase B
PAKt: phosphorylated protein kinase B
OXPHOS: oxidative phosphorylation

## Acknowledgements

The authors wish to thank Sen Hou for technical assistance. This work was supported by grants from the Heart and Stroke Foundation of Canada, NSERC, CHRIM (to G.M.H). G.M.H. is the Canada Research Chair in Molecular Cardiolipin Metabolism.

## Author contributions

H. Zegallai, E. Abu-El-Rub, E. Mejia, J. Field and L. Cole performed all experiments and analyzed the data. J. Gordon provided reagents and analytic tools. A. Marshall, H. Zegalli and G. Hatch were responsible for conceptual design. H. Zegallai and G. Hatch wrote the manuscript. All authors edited the manuscript.

